# The predictive structure of navigation determines mouse CA1 representational structure in a multicompartment environment

**DOI:** 10.1101/2024.11.25.625265

**Authors:** Sergio A. Pecirno, Alexandra T. Keinath

**Affiliations:** Department of Psychology, University of Chicago Illinois, 1007 W Harrison St., Chicago, IL, 60607, USA

## Abstract

Predictive theories of cognitive mapping propose that these representations encode the predictive relationships among contents as experienced by the navigator. One hallmark of these theories is that representational structure in complex environments can be predicted from the behavioral history of the navigator. Here, we image neural activity in hippocampal CA1 as initially naïve mice repeatedly navigate a multicompartment environment to test whether representational structure in this subregion is consistent with these predictions. We find that different mice instantiate different patterns of remapping across identically shaped compartments. Within mouse, compartments with more similar predictive navigational histories on a particular spatiotemporal scale are represented more similarly, accounting for these individual differences. Manipulating navigational options induces reorganization of the CA1 structure which specifically resembles the new predictive navigational structure on this scale. Through computational modeling we show that a combination of predictive encoding and geometrically structured inputs can uniquely account for this pattern of results. Together, these results demonstrate that the structure of CA1 representations in complex environments can be predicted from the behavioral history of the navigator, consistent with predictive theories of cognitive mapping.

---

In order to survive and thrive, organisms from mice to humans rely on mnemonic representations of the external world and their relationships to it^1^. Referred to as *cognitive maps*, these representations are instantiated by the coordinated activity of spatially tuned neural populations in the hippocampus and neighboring cortices^2,3^. Classically, cognitive maps are thought to represent geometric relationships – distances and/or angles – among their contents, motivated by the plethora of functional cell types tuned to geometric features throughout the hippocampal formation and neighboring cortices^2,4–18^. However, recent theories posit instead that cognitive maps represent the predictive relationships among their contents as experienced by the navigator^19–22^, with geometrically structured inputs providing a scaffold upon which predictive structure can be learned^3,23–25^.

Because the geometric layout of an environment often determines the predictive relationships that the navigator experiences, and modifying navigation often entails modifying the layout of the world, the outcomes of traditional and predictive models can be difficult to disentangle. Nevertheless, there are fundamental differences between these theories. One hallmark of predictive theories is that the way in which a navigator explores a space is a critical determinant of the structure of their cognitive map^19^. If navigators explore the same space differently, then their cognitive maps should be structured differently in a way which can be predicted from their navigational patterns. These differences should manifest across levels of explanation from neural instantiations through behavior, from the activity of single cells through the representational geometry of neural populations.

Motivated in part by these predictions, a growing body of work is contrasting behavioral and geometric determinants of neural activity in the hippocampus and functionally related cortices^6,13,26–34^. In many cases this work has demonstrated behavioral modulation of activity at the level of single cells and moment-to-moment population trajectories which is roughly consistent with predictive theories^13,29,30,33–35^ and which challenges traditional theories. In addition to these predictions, predictive theories also make concrete claims about how behavior should determine the representational similarity among locations in complex environments consisting of multiple subspaces^19^. Prior work has demonstrated that representational similarity in the hippocampus – or its converse, *remapping* – depends on the features shared between spaces^*36–40*^, past experience with those spaces^41–45^, and the broader reference frames in which those spaces are situated^43^. Individual differences in the degree of remapping within the same paradigm have been observed^46^, suggesting that remapping is not determined by world features and prior experience alone and can vary between navigators. Each of these characteristics can be understood within a predictive framework. However, more specific tests of whether patterns of hippocampal remapping in complex spaces conform to a predictive framework have failed^19,47^, calling into question whether the structure of hippocampal representations in such spaces can be understood through this lens.

Notably, this previous test included multiple complexities – such as extensive prior experience in the environment and short-term changes to navigational options and the reward landscape – which make deriving concrete predictions from a predictive framework more challenging. Here, we test whether patterns of hippocampal remapping in complex spaces conform to a predictive framework in a simpler paradigm which avoids some of these complexities. Specifically, we characterize population activity in hippocampal CA1 as initially naïve mice repeatedly explore four identically shaped connected compartments in the absence of reward. To do so with high fidelity and across long timescales, we rely on chronic miniscope imaging. We test three hypotheses. If CA1 representational structure conforms to a predictive framework, then (i) mice that navigate differently should exhibit different patterns of remapping across compartments, (ii) within mouse, compartments with more similar predictive navigational histories should be represented more similarly, and (iii) changing the navigational trajectory of a mouse should change its representational structure to better resemble the new predictive structure. Evidence against any of these hypotheses would support an alternative theory.

In all cases our data support the predictive claims. That is, we find that different mice exhibit different patterns of remapping across compartments. These patterns coincide with the similarity of predictive navigational histories on a seconds-long timescale, modestly overweighting early experience. Manipulating the navigational trajectory of the mouse induces a reorganization of the CA1 representational structure which specifically resembles the new predictive structure on this timescale, with a reversion to the previous pattern when relaxed. Leveraging computational modeling, we show that key results can be accounted for by a combination of predictive encoding operating on geometrically structured inputs; neither predictive encoding nor geometrically structured inputs were on their own sufficient to reproduce our pattern of results. Together, these results demonstrate that the structure of hippocampal representations in complex environments can be predicted from the behavioral history of the navigator, consistent with predictive theories of cognitive mapping.

## Results

### Characterizing CA1 representational structure in a multicompartment environment

We recorded daily from hippocampal CA1 via miniscope calcium imaging (Fig. 1a) as initially naïve mice (n = 11; 6 male) navigated a radial multicompartment environment. This environment consisted of four 20 cm by 35 cm rectangular compartments connected by a central chamber (Fig. 1b). Each session lasted either 30 or 40 min depending on imaging signal strength to maximize the amount of data while minimizing the impact of photobleaching. In the first phase of the experiment, initially naïve mice navigated this environment freely for between five and fifteen sessions (n = 121 sessions total). We refer to these sessions as *free*_*1*_. The environment was not baited with rewards during any phase of the experiment.

**Figure 1.**
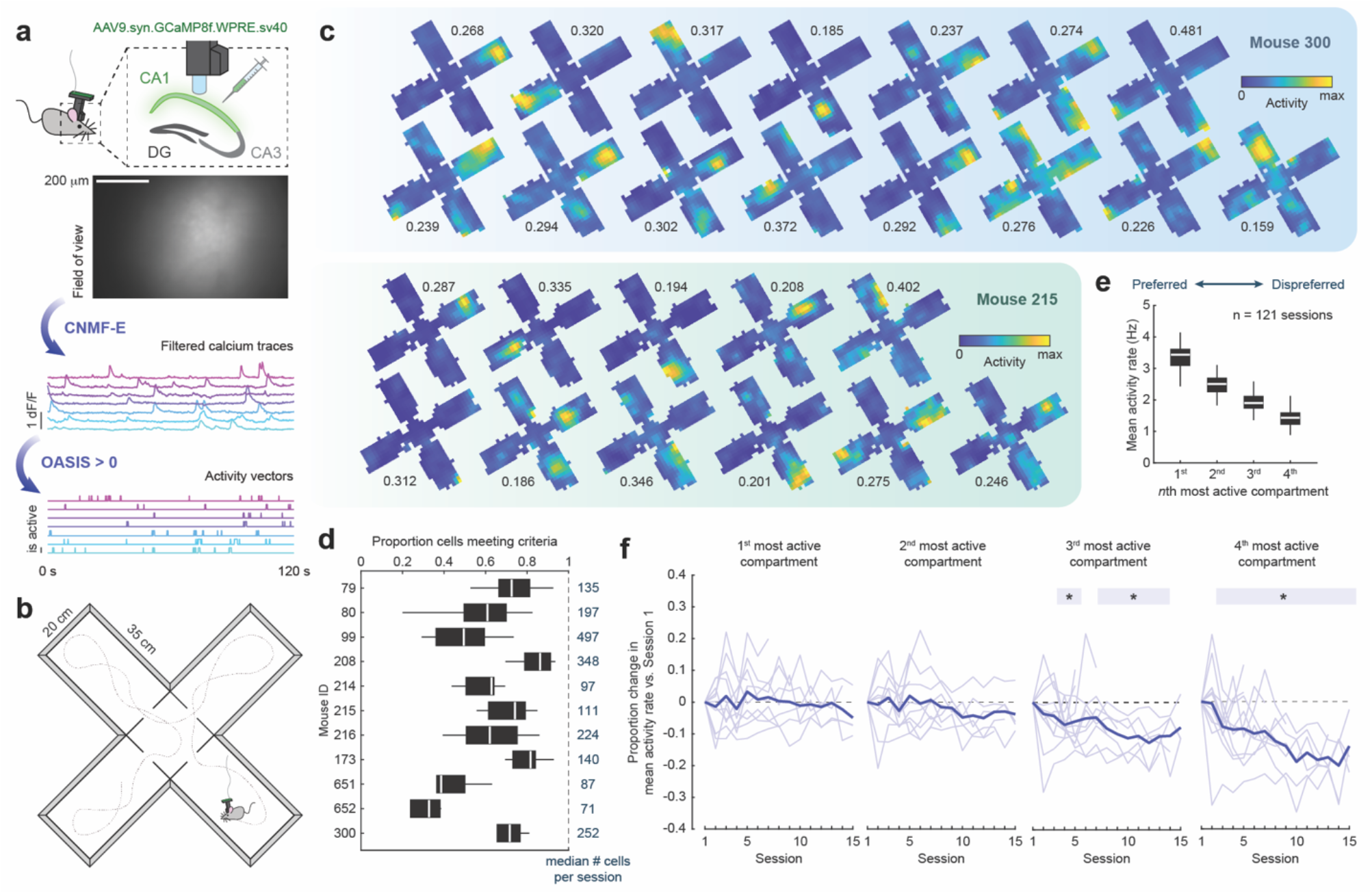
Characterizing mouse CA1 in a multicompartment environment via miniscope imaging. (a) Schematic of miniscope imaging and data processing pipeline. (b) Schematic of the multicompartment environment. (c) Example rate maps from simultaneously-recorded cells for one session from two mice. Rate maps normalized from zero (blue) to peak activity rate (yellow). Peak activity rate in proportion of active frames noted for each map. (d) Proportion of cells with split-half stability and spatial information content exceeding the 95^th^ percentile of their shuffled controls across sessions for each mouse. Median number of cells per session for each mouse noted on the right. (e) Mean activity rate as a function of most active to least active compartments. (f) Change in mean event rate versus the first session, as a function of most active to least active compartments. Soft shaded lines are individual mice. Bold line is the mean across mice. Shaded box denotes sessions for which p < 0.05. Significance markers denote the outcome of a t-test versus zero for a given compartment and session. *p<0.05

For each session, imaging data were first motion corrected^48^. Next, cells were segmented, and their calcium traces were extracted via constrained nonnegative matrix factorization (CNMFE; Fig. 1a)^49,50^. For each trace, the likelihood of spiking events giving rise to that trace were inferred through a second-order autoregressive deconvolution algorithm^51^ (Fig. 1a). This nonnegative vector was then binarized such that any frame with a nonzero likelihood was treated as active (i.e. 1), otherwise it was treated as not active (i.e. 0). All further analysis was conducted on this binary activity vector.

To characterize the extent to which spatial location modulated the activity of each cell, we computed a rate map for each cell summarizing the mean activity of that cell as a function of location within the environment (Fig. 1c). Next, we computed the split-half stability and spatial information content of this rate map. To assess the significance of these measures, we compared these values to surrogate distributions computed by circularly shifting the activity vector relative to the position vector 1000 times. Cells with split-half stability and spatial information content exceeding the 95^th^ percentiles of both distributions were considered spatially tuned and included in further analysis, though relaxing these criteria did not qualitatively change our results. Generally, the majority of cells met these criteria (Fig. 1d).

Prior work in multicompartment environments has demonstrated that individual CA1 cells tend to be active in multiple compartments^52,53^, making comparisons between compartments meaningful. We confirmed this was the case in our data. To do so, for each cell we computed its mean activity rate in each compartment and sorted these from the most active to least active compartments. Across sessions, we found that mean activity rates remained high in all compartments, with on average at least ∼1 active frame per second even in dispreferred compartments (Fig. 1e). The difference in activity rate between preferred and dispreferred compartments was approximately 3-fold. Interestingly, we also observed a change in mean activity rates with experience in dispreferred (but not preferred) compartments (Fig. 1f), consistent with the development of sparsity and/or sharpening of the spatial code across experience^54^.

### Different mice instantiate distinct and persistent patterns of remapping across identical compartments

Predictive theories claim that, all else being equal, CA1 representational structure should be determined by the predictive structure of one’s navigational trajectory. If so, then we would expect mice in our paradigm to instantiate different patterns of remapping between compartments insofar as they navigate with different trajectories. On the other hand, purely geometric theories might predict that remapping is dictated by the geometric features of the environment, such as the distance between compartments or the relative orientations of the compartments^52,53^. If so, then we would expect mice in our paradigm to instantiate similar patterns of remapping, perhaps corresponding to key geometric features.

To address these possibilities, we characterized the degree of remapping between compartments in our data using a population vector approach. To this end, for each cell we first computed a separate rate map for each compartment (Fig. 2a). Next, for each session we computed the population vector (PV) correlations between all pairwise comparisons of compartment rate maps, aligned by their entryways and excluding unsampled pixels (Fig. 2a). PV correlations summarize the degree to which two spaces are represented similarly at the level of the neural population in a way that captures changes in both firing rates and spatial tuning. Computing all pairwise PV correlations between compartments yields a 4 x 4 matrix which we refer to as the *PV structure* (Fig. 2a). Finally, we correlated PV structures between sessions and compared correlations between different sessions from the same mice to correlations between sessions from different mice (Fig. 2b).

**Figure 2.**
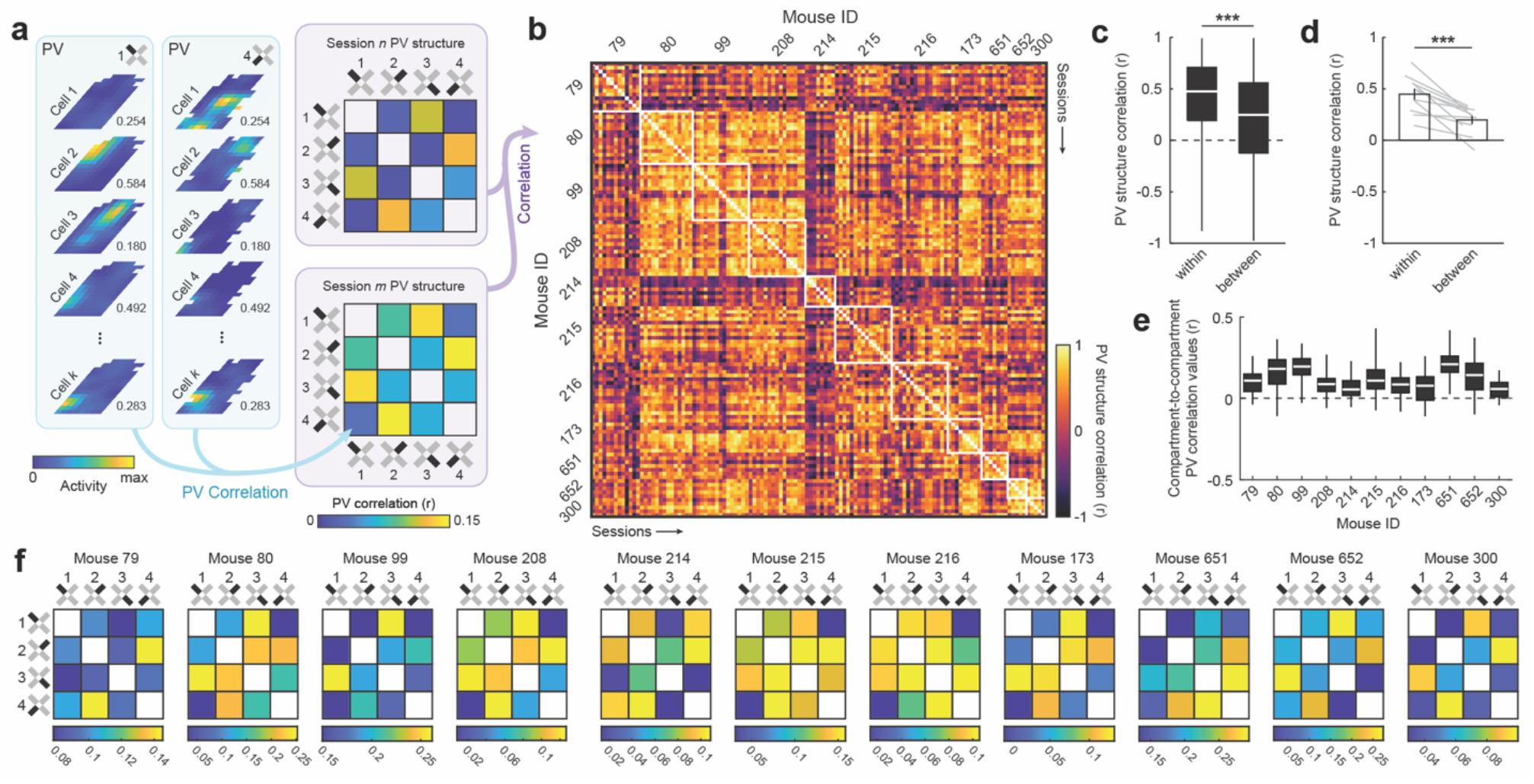
Individual differences in patterns of remapping across compartments in a multicompartment environment. (a) Schematic of PV structure analysis pipeline. Rate maps scaled to peak activity rate across all compartments. This peak rate noted next to each map. (b) PV structure correlations across all pairwise comparisons of sessions. White boxes bound session comparisons from the same mice. (c) PV structure correlations for comparisons within mouse versus between different mice, treating each pairwise session comparison independently (rank-sum test: z(3143353) = 11.879, p = 1.513e-32). (d) PV structure correlations for comparisons within mouse versus between mice, treating each mouse as the unit of analysis (signed-rank test: p = 9.766e-04). Soft lines denote individual mice. (e) Distribution of compartment-to-compartment PV correlation values for each mouse. (f) PV structures averaged across sessions for each mouse. ***p<0.001

Consistent with the predictive theory, comparisons of sessions from the same mouse resulted in significantly higher correlations than comparisons between mice (Fig. 2c,d). Moreover, despite the limited number of comparisons defining the PV structure (i.e. six pairwise comparisons between the four compartments), these differences were so reliable that a simple Euclidean distance based classifier could correctly predict mouse identity well above chance (accuracy = 45.45%, chance = 9.09%, p = ∼0; binomial test). This reliable structure was present even though PV correlation values between compartments were generally low in magnitude (Fig. 2e), suggesting that even weak correlations in representational structure can be stable and informative^55^. Even averaging within mouse, PV structures varied widely between mice and qualitatively did not come to resemble a consistent geometric feature of the compartmental layout (i.e. distance, entryway angle, long axis, etc.; Fig. 2f). Similar results were observed when correlating mean activity rates as the measure of remapping (Fig. S1), indicating that our results are not dependent upon the rate map-based assumptions underlying PV correlations. Together, these results demonstrate that mice instantiate idiosyncratic patterns of remapping across compartments in this multicompartment paradigm.

### Idiosyncratic CA1 representational structure matches the predictive structure of navigation on a particular spatiotemporal scale

Our previous results demonstrate reliable individual differences in CA1 representational structure in mice navigating our multicompartment environment. While these differences could arise from predictive cognitive mapping, individual differences are not on their own unique to this theory. For example, a geometric theory might predict individual differences if navigators are attending to or naturally emphasize different features of the world. Therefore, we next tested the more specific claim of predictive theories: that individual differences in CA1 representational structure arise due to idiosyncratic navigational differences. If correct, then compartments with more similar predictive navigational structures should be represented more similarly in CA1.

To test this possibility, we leveraged one formalization of predictive cognitive mapping, the successor representation^19^. At a high level, the successor representation defines the behavioral determinants of hippocampal representational structure as the temporally discounted probabilities of transitioning between locations within an environment. This is expressed as a successor matrix *M* such that

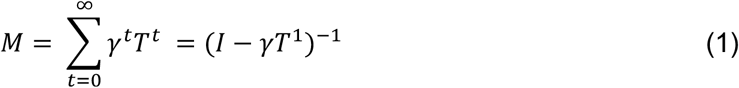

where *T*^*n*^ is the *n*-step transition matrix capturing the likelihood of transitioning from one location to another after *n* timesteps, *γ* is the temporal discount factor which determines how heavily transitions near in time versus far in time are discounted, and *I* is the identity matrix.

In our case, for a given session *s* we can compute the successor matrix *M*_*s,k*_ separately for each compartment *k*. Correlating these matrices between all pairwise comparisons of compartments yields a 4 x 4 matrix which we can compare to the PV structure for that session. If the predictive claim is correct, then these matrices should be correlated. Importantly though, in our paradigm mice explored the environment for multiple sessions, and it is not obvious whether navigation during every session should have an equal impact on CA1 representational structure. While this is certainly a possibility, it is also possible that more recent behavior or early behavior has an exaggerated impact on determining CA1 representational structure. The longitudinal nature of our recording paradigm allows us to test these possibilities. To do so, for each session *s* and compartment *k* we computed a cumulative one-step transition matrix *C*_*s,k*_ by including all navigational data up to and including that session such that

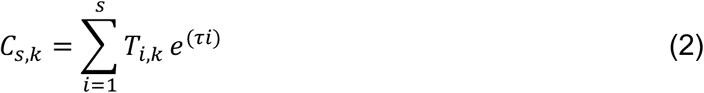

where *T*_*i,k*_ is the one-step transition matrix for session *i* and compartment *k* and *τ* determines the exponential weight with which each session contributes to the cumulative transition matrix. A negative value of *τ* will overweight early session behavior. A positive value of *τ* will overweight more recent behavior. A *τ* of zero will equally weight data from all sessions. Substituting our weighted cumulative one-step transition matrix for *T*^1^ in equation (1), we can compute the cumulative successor matrix *M*_*s,k*_ for each session *s* and compartment *k* such that

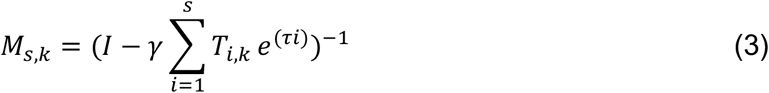

Finally, we can correlate *M*_*s,k*_ between all pairwise comparisons of compartments (excluding unsampled pixels), yielding the 4 x 4 matrix which we refer to as the *cumulative successor structure* (Fig. 3a). This matrix summarizes the similarity in predictive navigational structure between compartments dependent upon two parameters, *γ* and *τ*. These parameters determine the time horizon of the encoded predictive structure and the weighting of cumulative experience, respectively.

**Figure 3.**
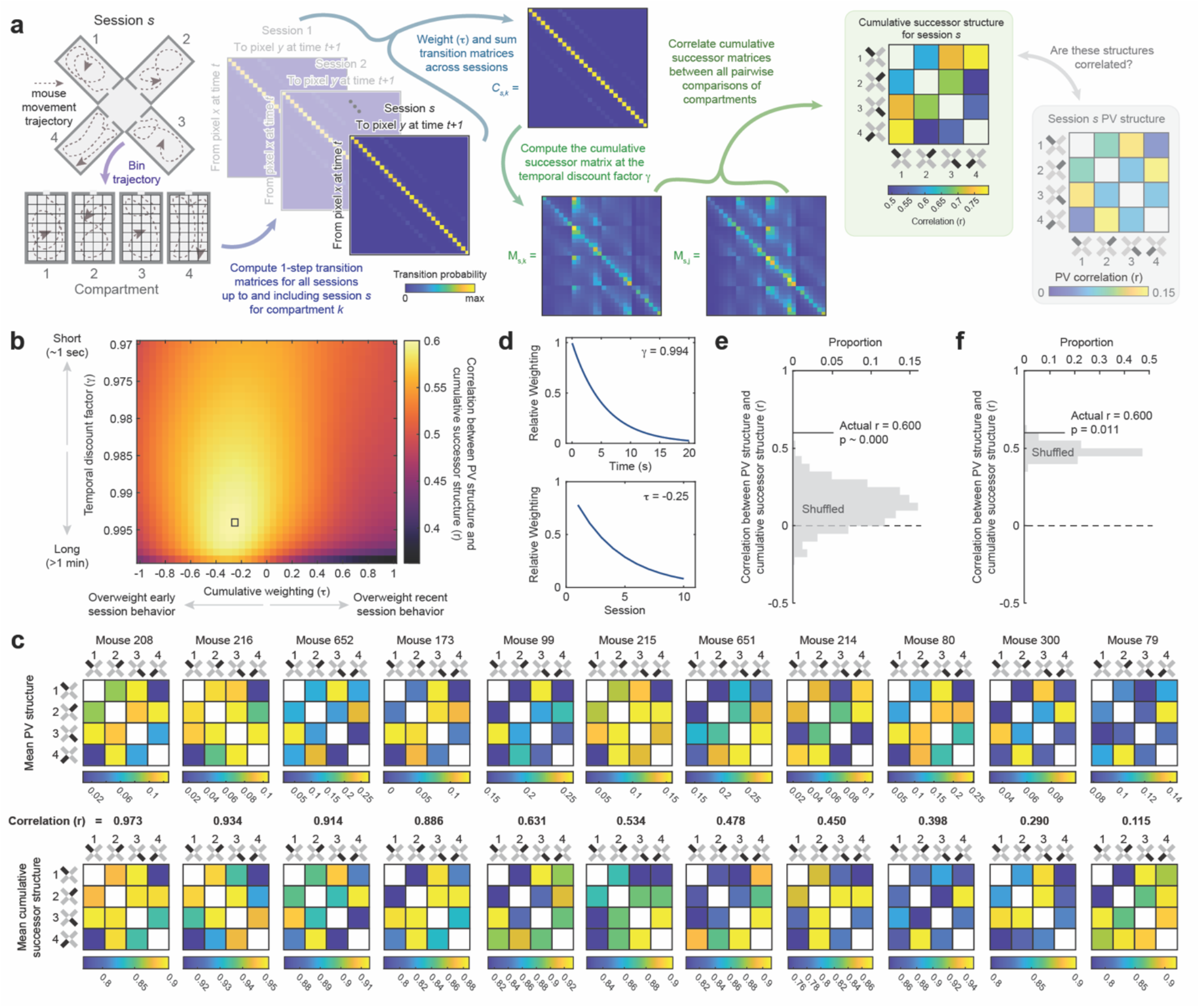
CA1 PV structure matches the similarity of predictive navigational structure on a particular spatiotemporal scale. (a) Schematic of pipeline for computing cumulative successor structure. (b) Grid search where γ and τ are varied and the mean cumulative successor structure is correlated with the mean PV structure for each mouse. (c) Mean PV structures and mean cumulative successor structures for all mice with correlations noted. Ordered from most to least correlated. (d) Relative weightings derived from γ and τ at maximizing parameterizations. (e) Correlation between PV structure and cumulative successor structure against a shuffled control where transition probabilities are shuffled across compartments, mice, and sessions and controlling for the grid search. (f) Correlation between PV structure and cumulative successor structure against a shuffled control where transition probabilities are shuffled across sessions within mouse and controlling for the grid search.

To determine whether cumulative successor structures match CA1 PV structures at any parameterization, we conducted a grid search where *γ* and *τ* were varied and the mean successor structure was correlated with the mean PV structure for each mouse. This analysis revealed that mean successor structures were highly correlated with mean PV structures at a particular subset of parameterizations, with a maximum at *γ* = 0.994 and *τ* = -0.25 (Fig. 3b,c; Fig. S2a). These parameters correspond to a seconds-long timescale for encoding predictive structure, modestly overweighting early experience (Fig. 3d). To assess the significance of the high correlation we observed at this parameterization (mean r = 0.600), we considered several controls. First, we compared this value to the distribution we might expect from generic mouse behavior by randomly shuffling transition probabilities (i.e. *T* in Equation 3) between compartments, sessions, and mice prior to computing cumulative successor structures. To account for the selection of the maximizing parameterization from our grid search, we repeated our grid search for each shuffle and took the maximum correlation for that shuffle. The true correlation exceeded this control (nonparametric p ∼ 0.000, 1,000 shuffles; Fig. 3e). Next, we tested whether the specific order of experience mattered by randomly shuffling navigational trajectories across sessions separately for each mouse before computing cumulative successor structures and choosing the maximum correlation across the grid search for each shuffle. Again, the true correlation exceeded this control (nonparametric p = 0.011, 1,000 shuffles; Fig. 3f). Repeating these analyses but treating individual sessions as the unit of analysis rather than individual mice yielded similar results (Fig. S2). Substituting mean activity rates as the measure of remapping also yielded similar results (Fig. S2), indicating that the concordance between predictive structure and degree of remapping we observe is not dependent upon the rate map-based assumptions inherent in PV correlations. We note that positive values of *τ* which overweight current session experience yielded numerically weaker correlations between successor structures and PV structures. This provides additional evidence that the concordance we observe between these structures at maximizing parameterizations is not due to low-level within-session confounds. Together, these results demonstrate that compartments with more similar predictive navigational structures on a particular spatiotemporal scale are represented more similarly in CA1, consistent with a predictive theory of cognitive mapping.

To gain more insight into how predictive navigational structure varied within and between mice on this critical timescale (*γ* = 0.994), we next computed individual session (non-cumulative) successor structures and correlated these structures between sessions (Fig. 4a). Successor structures were modestly more correlated within mouse than between mice (Fig. 4b,c), suggesting some consistency in behavioral patterns across sessions within mouse. However, this varied considerably between mice, with some mice showing little to no consistency in successor structures across sessions. This variability is useful for distinguishing the relative weightings of cumulative behavior (i.e. *τ*), as consistent predictive structure across sessions makes it difficult to parse out the relative contributions of individual sessions. It also suggests that the greater degree of consistency we observe in the CA1 PV structure across sessions is not due to consistent behavior from session to session alone but rather reflects integration of experience across time.

**Figure 4.**
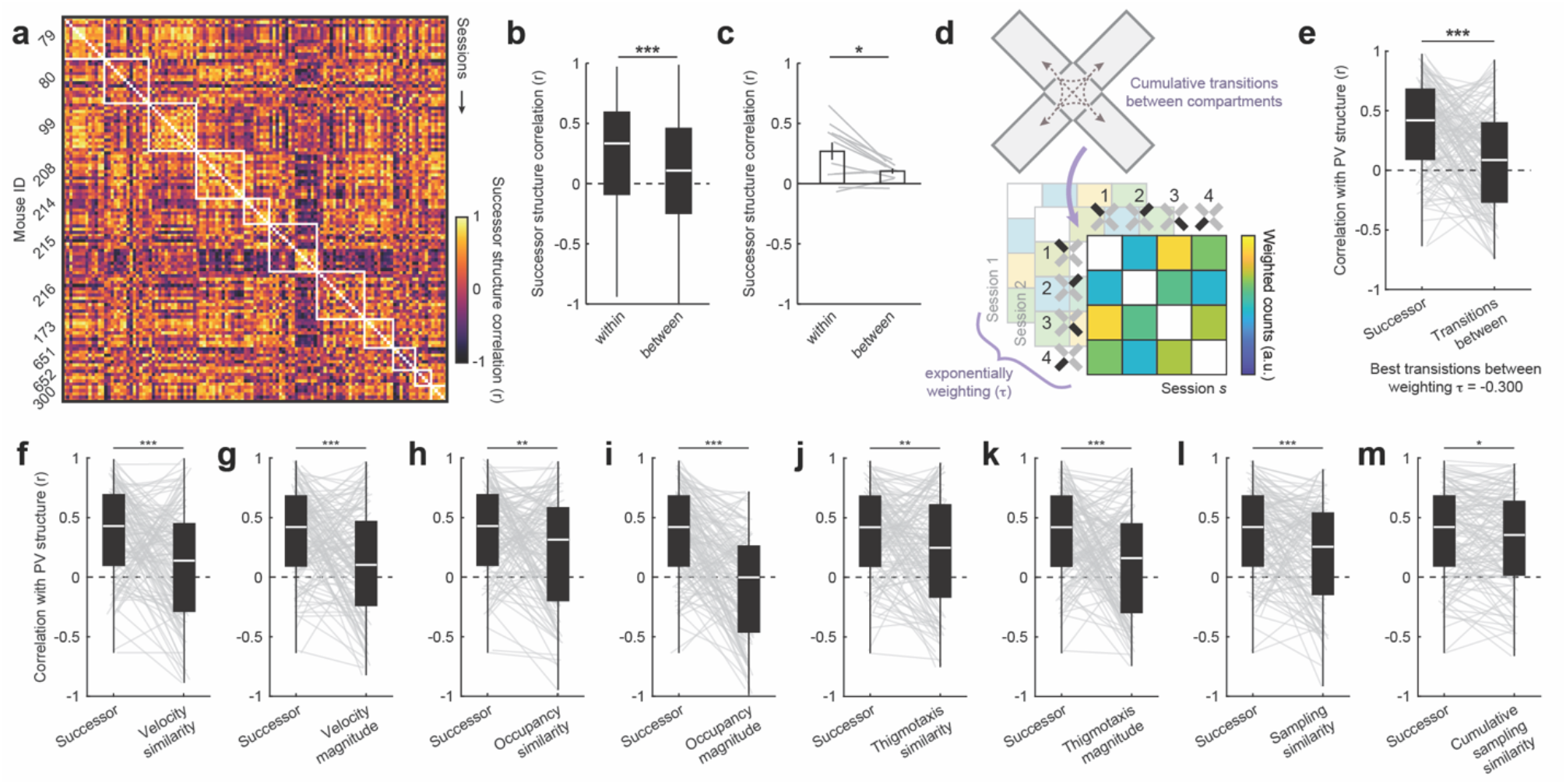
Successor structures are modestly stable across sessions and cumulative successor structures better explains PV structure than other behavioral variables. (a) Noncumulative successor structure correlations across all pairwise comparisons of sessions. White boxes bound session comparisons from the same mice. Note that some mice exhibit high correlations across sessions while others do not. (b) Successor structure correlations for comparisons within mouse versus between different mice, treating each pairwise session comparison independently (rank sum test: Z(2964100) = 8.466, p = 2.545e-17). (c) Successor structure correlations for comparisons within mouse versus between different mice, treating each mouse as the unit of analysis (sign rank test: p = 3.223e-02). (d) Schematic for computing cumulative weighted transitions between compartments. (e) PV structures were more highly correlated with cumulative successor structures than cumulative transitions between compartments (sign rank test: Z(5609) = 4.962, p = 6.963e-07). Sessions treated independently. (f) As in (e), except for velocity similarity (Z(5405) = 4.435, p = 9.219e-06). (g) As in (e), except for higher mean velocities (Z(5253) = 4.042, p = 5.309e-05). (h) As in (e), except for the similarity of occupancy times (Z(4769) = 2.790, p = 5.276e-03). (i) As in (e), except for higher mean occupancy times (j) As in (e), except for more similar degrees of thigmotaxis (Z(4941) = 3.235, p = 1.218e-03). (k) As in (e), except for higher mean degrees of thigmotaxis (Z(5285) = 4.124, p = 3.718e-05). (l) As in (e), except for the similarity of sampling distributions (Z(5270) = 4.086, p = 4.397e-05). (m) As in (e), except for the similarity of cumulative sampling distributions (Z(4528) = 2.166, p = 3.029e-02). *p<0.05, **p<0.01, ***p<0.001

Finally, we asked whether other behavioral determinants might explain CA1 PV structure equally well. We first tested whether transitions between compartments were equally correlated with PV structure, as might be expected from some instantiations of a predictive theory^19,47^. To this end, for each session we computed the cumulative number of transitions between each pair of compartments across all sessions up to and including that session (Fig. 4d). As before, it is unclear whether experience during different sessions should be weighted equally. Thus, we weighted the contributions of each session as in Equation (2) and determined the maximizing *τ* through a grid search at the same resolution as before (i.e. Fig. 3b). Even at maximizing *τ*, these matrices were significantly less correlated with PV structures than cumulative successor structures (Fig. 4e).

Next, we asked whether differences in velocity, time spent in each compartment, thigmotaxis, or general sampling patterns could equally account for PV structure. Neither the similarity of velocity – measured as the inverse absolute difference between mean velocity in each compartment – nor higher mean velocities could account for PV structure as well as cumulative successor structure (Fig. 4f,g). Neither the similarity of occupancy times – measured as the inverse absolute difference between total occupancy time in each compartment – nor longer mean occupancy times could account for PV structure as well as cumulative successor structure (Fig. 4h,i). Neither similar degrees of thigmotaxis – measured as the inverse absolute difference between the proportion of time spent within 5cm of a wall in each compartment – nor higher mean degrees of thigmotaxis could account for PV structure as well as cumulative successor structure (Fig. 4j,k). Even the similarity of sampling patterns – measured as the correlation between pixel occupancy maps within each compartment – could not account for PV structure as well as cumulative successor structure. This was true regardless of whether we compared only the sampling for a given session or the cumulative sampling for all sessions up to and including that session (Fig. 4l,m). Together, these results demonstrate that cumulative successor structures uniquely resemble CA1 representational structure, even when compared to other associated behavioral metrics.

### Manipulating navigational options induces reorganization of CA1 representational structure to match the new predictive structure of navigation

Our previous results demonstrate that CA1 exhibits individual differences in the degree of remapping between compartments which coincides with the similarity of predictive structure on a particular spatiotemporal scale. While these results provide correlational evidence consistent with a predictive theory of cognitive mapping, these theories make a stronger causal claim. That is, changing the movements of a navigator should induce a corresponding change in representational structure to match the new predictive structure. We next sought to test this prediction. To this end, in four of our mice we followed the initial free navigation phase (15 sessions; *free*_*1*_) with an additional phase of limited navigation (7 sessions; *limited*) and finally a subsequent free navigation (21 sessions; *free*_*2*_; *Fig. S3*). During the limited phase, each session consisted of locking the mouse in each compartment for four minutes at a time, twice per session. Locking was achieved by the experimenter placing a small barrier in the entryway. This barrier was made of the same material as the walls and fit snuggly in the entryway, matching its dimensions. When each four-minute epoch ended, the door was removed and the mouse freely navigated in clockwise order to the next compartment.

Extensive evidence indicates that manipulating environmental features^37,38,43–45,56^ and behavioral contingencies ^57–59^ can induce remapping in hippocampal CA1. We therefore reasoned that limiting navigational options, a novel experience for the mouse, would lead to remapping in CA1 during which new predictive structure might be encoded. Moreover, although this manipulation does not enforce a particular change to their navigational trajectories, free navigation typically involved shorter bouts of exploration within each compartment (median bout duration = 11.13 ± 4.92 s; mean ± standard deviation across Free_1_ sessions). Thus, we suspected that limiting options would also lead mice to deviate from their free navigational patterns.

We first confirmed that our manipulation did indeed induce remapping in CA1. To do so, we correlated PV structures across all sessions within each mouse and compared correlations between different phases of this experiment. Limiting navigational options led to a clear reorganization of PV structure on averaged and separately in each mouse (Fig. 5a). Quantifying this, correlations between free phases and the limited phase were significantly lower than correlations within the same phase (rank sum tests: Z(1891217) = 19.020, p = 1.163e-80) or between free_1_-free_2_ (Z(926848) = -13.985, p = 1.917e-44; Fig. 5b). Correlations between free_1_-free_2_ were numerically high but significantly weaker than same phase correlations (Z(1874165) = 6.447, p = 1.143e-10). This pattern of results indicates rapid remapping during the limited manipulation, followed by a near return to the initial representational structure when this manipulation was relaxed.

**Figure 5.**
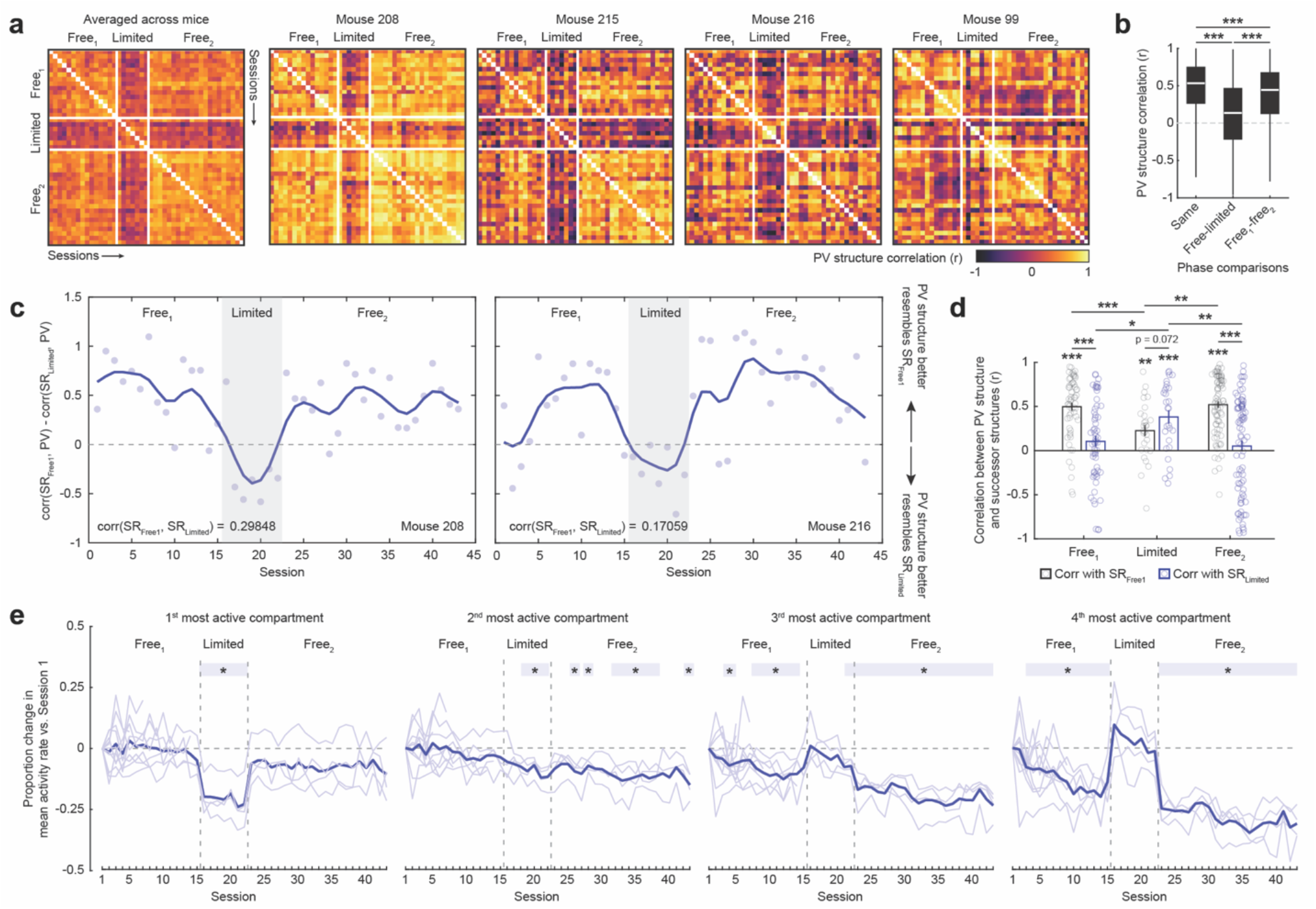
Limiting navigational options induces remapping which encodes the new predictive structure. (a) PV structure correlations across all 43 sessions averaged across mice and separately for each mouse. Experimental phase boundaries denoted by white lines. (b) PV structure correlations for comparisons between sessions from the same phase, between free and limited phases, and between Free_1_ and Free_2_ phases. (c) Difference in the correlation between PV structure and SR_Free1_ and SR_Limited_ for two example mice. Dots indicate individual session values. Line depicts these data smoothed with a gaussian kernel with 1.5 session standard deviation. Correlation between the SR structures denoted in the lower right for each mouse. (d) Correlation between PV structure and both SR_Free1_ and SR_Limited_ for all sessions, aggregating across mice. For complete statistical information, see Supplementary Table 1. (e) Change in mean event rate versus the first session, as a function of most active to least active compartments. Soft shaded lines are individual mice. Bold line is the mean across mice. Shaded box denotes sessions for which p < 0.05. Significance markers denote the outcome of a t-test versus zero for a given compartment and session. *p<0.05, **p<0.01, ***p<0.001

Given that we observed rapid remapping during the limited phase, we next tested the key hypothesis that patterns of remapping during the limited phase would *specifically resemble the new predictive structure*. To do so, for each mouse we computed the mean cumulative successor structure for the free_1_ and limited navigation phases at the critical spatiotemporal scale determined above (*γ* = 0.994 and *τ* = -0.25). We refer to these mean cumulative successor structures as SR_Free1_ and SR_limited_. These structures were only partially correlated with one another within each mouse (r_99_ = 0.699, r_208_=0.299, r_215_ = -0.330, and r_216_ = 0.171). We then correlated each of these successor structures with the PV structure for each session and compared these correlations across phases (Fig. 5c,d). This analysis revealed that during the free_1_ phase, PV structures were highly correlated with SR_Free1_ but uncorrelated with SR_limited_. Crucially, during the limited phase PV structures became correlated with SR_limited_, representing a significant increase over free_1_ phase correlations. Correlations with SR_limited_ numerically exceeded those with SR_Free1_, though this difference was statistically marginal. Finally, during free_2_ the pattern of correlations reverted to that of free_1_. These results did not depend on the inclusion of pixels near the entryway in the analysis (Fig. S4). Together, these results demonstrate that our manipulation specifically induced a pattern of remapping which matched the new predictive structure at the critical spatiotemporal scale, consistent with a predictive framework.

Our findings indicate that limiting navigational options led to rapid encoding of new predictive structure while the subsequent return to free navigation prompted a return of the free_1_ representational structure. This pattern suggests that the rate of encoding of predictive structure appears to be specific to the experimental phase. Though speculative, we asked whether we could observe other hallmarks of encoding which might corroborate this possibility. We noted that during free_1_ activity tended to become sparser over repeated sessions, as reflected by a decrease in the mean activity rate in dispreferred compartments (Fig. 1f). This decrease roughly matched the optimal weighting of predictive structure across sessions (*τ* = -0.25). We therefore characterized sparsity across the entire experiment utilizing our full dataset (n = 233 sessions). This analysis revealed that during limited navigation mean activity rates in dispreferred compartments returned to their initial levels but immediately reverted to their pre-limited rates during the subsequent free_2_ (Fig. 5e). Qualitatively, during the limited phase mean activity rates in dispreferred compartments decreased across sessions in a way echoing free_1_, while free_2_ continued the trending decrease from free_1_. Mean activity rates in the most active compartment also decreased significantly during the limited phase and returned to their free_1_ levels during the subsequent free_2_ phase. Overall, this pattern provides tentative evidence of phase-specific activity changes coinciding with enhanced encoding of predictive structure. Moreover, these findings suggest that the sparsity of the CA1 representation might serve as a proxy for the rate at which predictive structure is encoded.

### A computational model encoding predictive structure over geometrically structured inputs uniquely accounts for key results

We have shown that in a complex environment CA1 instantiates idiosyncratic patterns of remapping between compartments which are correlated with the similarity of predictive navigational structure on a particular spatiotemporal scale. To gain insight into how this pattern of results might arise, we implemented a series of computational models intended to mimic the input-output transformation carried out by CA1. We varied two aspects of these models: the spatial structure of their inputs and the transformation they compute on these inputs.

We considered inputs with two types of spatial structure. In both cases, inputs consisted of cells tuned to a single preferred location in each compartment. In the case of *geometrically structured inputs*, the preferred locations were in geometrically identical locations in each compartment for a given cell (Fig 6a). In the case of *randomly structured inputs*, the preferred location randomly differed between each compartment for a given cell (Fig 6a). Contrasting the two will help us infer whether spatial tuning of inputs alone is sufficient to reproduce our results, or whether geometric structure plays an important role.

**Figure 6.**
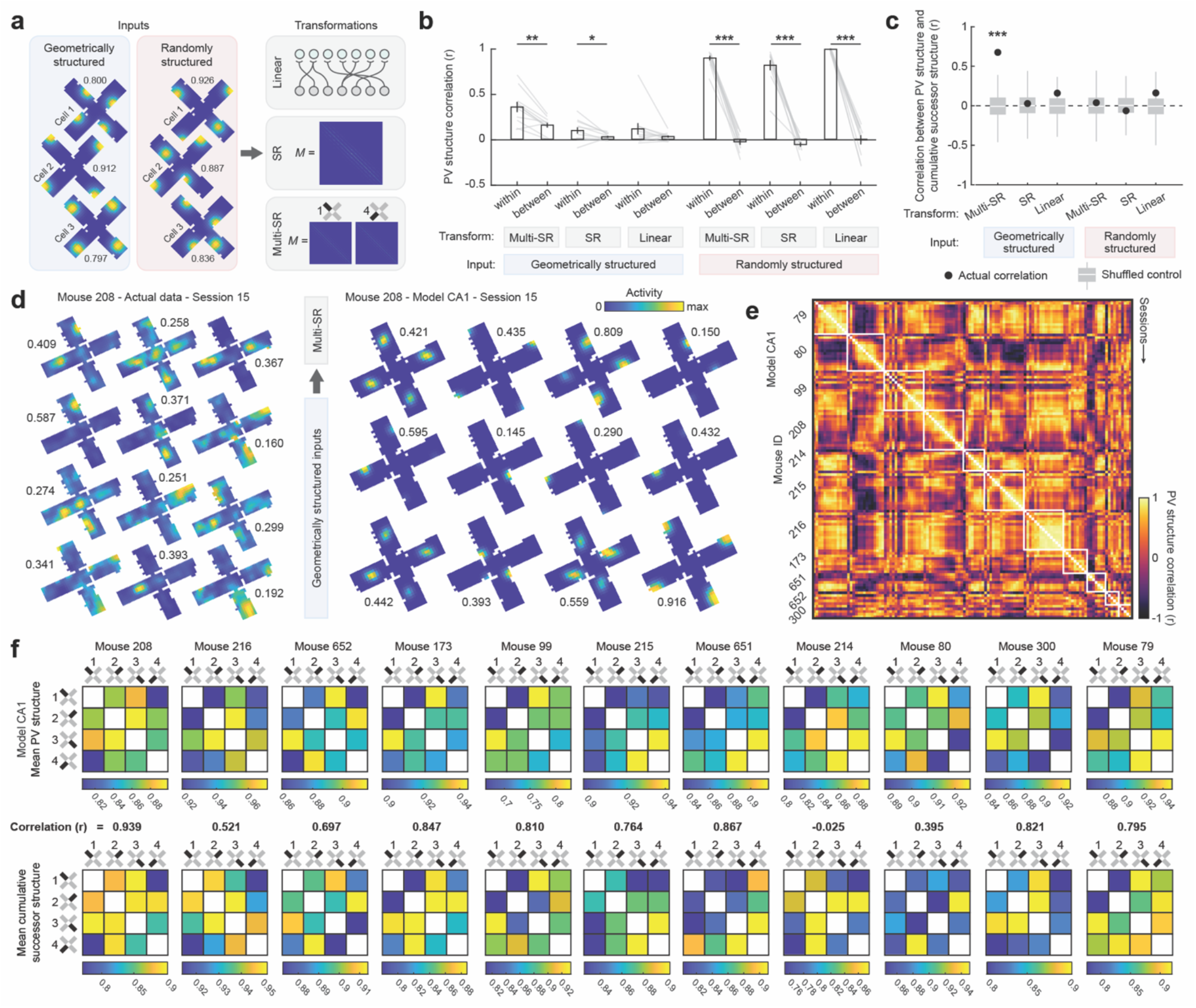
A predictive model with geometrically structured inputs replicates key results. (a) Schematic depicting possible model configurations. Peak activity rate noted for each rate map. (b) PV structure correlations among sessions separately for comparisons within the same mouse or between different mice, treating each mouse as the unit of analysis and separately for each model (signed-rank tests; geo-multiSR, p = 2.930-03, geo-SR, p = 0.024, geo-linear, p = 0.101; all random input models, ps = 9.766e-04). Soft lines denote individual mice. (c) Correlations between the mean PV structure and the mean cumulative successor for each mouse, separately for each model. Black dots denote the true correlation. Grey box-and-whiskers denote the control distribution computed by shuffling cumulative successor structures across sessions (1,000 shuffles, nonparametric p-values estimated against surrogate distributions: geo-multiSR, p ∼ 0.000, geo-SR, p = 0.430, geo-linear, p = 0.065, rand-multiSR, p = 0.389, rand-SR, p = 0.690, rand-linear, p = 0.504). (d) Example rate maps for actual data and geo-multiSR model CA1. Peak activity rate noted for each rate map. (e) PV structure correlations across all pairwise comparisons of sessions for geo-multiSR model CA1. White boxes bound session comparisons from the same mice. (f) Mean PV structures for geo-multiSR model CA1 and mean cumulative successor structures for all mice with correlations noted, ordered as in Fig. 3c. *p<0.01 **p<0.01, ***p<0.001

We also considered three types of transformations. Firstly, we considered a simple linear transformation (*linear*), where each CA1 cell received a small number of equally weighted inputs (see Methods). Secondly, we considered a predictive transformation (*SR*) where CA1 learns an approximation of the successor representation over its inputs, as explored previously^24^. In this case, the weight matrix *M* which determines the strength of the connection from each input cell to each CA1 cell is an online approximation of the successor representation updated at each timestep according to

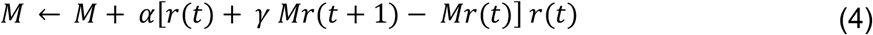

where *r*(*t*) is the vector of input firing rates at time *t, γ* is the temporal discount factor, and *α* denotes the learning rate. The total input *F*_*i*_ to each CA1 cell *i* is then determined by

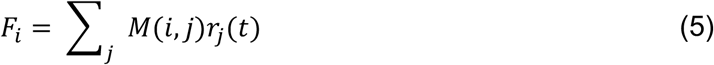

where *M* is the weight matrix at the current timestep computed in equation (4), and *r*_*j*_(*t*) is the firing rate of the *j*th input at the current timestep *t*. Thirdly, we considered a version of the predictive transformation where a separate successor representation *M* is learned for each compartment (*multi-SR*), and *F*_*i*_ for each CA1 cell is computed based on the *M* corresponding to the currently occupied compartment. Finally, for all transformations the output of each CA1 cell was thresholded such that only the top 10% most excited cells across the population were active during each timestep and normalized to range between zero and one (see Methods).

To carry out our simulations, we used the actual navigational trajectories of our mice from the Free_1_ phase of our experiment. For SR and multi-SR simulations, we used *γ* = 0.994 corresponding to the maximizing parameterization estimated above. To capture the decreasing weighting of cumulative experience over repeated sessions (i.e. *τ* = -0.25), the learning rate *α* was set to decay exponentially at *τ* = -0.25 from its initial value on session one (initial *α* = 0.005).

For each combination of input structures and transformations we first asked whether we observed reliable individual differences in representational structure, as we observed in our actual data. To this end, for each model we correlated PV structures between all pairwise comparisons of sessions and contrasted comparisons from within the same mouse versus between different mice (Fig. 6b). We found that all models with randomly structured inputs (which differed from mouse to mouse) recapitulated strong individual differences as indicated by higher correlations for comparisons within mouse than between different mice. On the other hand, geometrically structured inputs (which were identical across mice) did not recapitulate individual differences in every case. Only when these inputs were coupled with learning of predictive structure (i.e. an SR or multi-SR transformation) were significant individual differences observed. These differences were particularly strong when predictive structure was learned separately for each compartment (i.e. multi-SR). These results demonstrate that individual differences are consistent with predictive encoding of geometrically structured inputs, although they may alternatively reflect other between-mouse differences such as randomness in input structure.

Next, for each combination of input structures and transformations we asked whether model PV structures were correlated with cumulative successor structures, as we observed in our actual data. Computing cumulative successor structures at the maximizing parameters (*γ* = 0.994, *τ* = -0.25), we found that PV structures were significantly correlated with cumulative successor structures in only one case: when inputs were geometrically structured and predictive encoding was learned separately for each compartment (multi-SR). Neither factor was on its own sufficient, as other models with geometrically structured inputs or predictive encoding learned separately for each compartment failed to produce significant correlations. Interestingly, environment-wide encoding of predictive structure (SR) failed to produce significant correlations even when inputs were geometrically structured. This is because this model encodes only a single predictive structure for the entire environment, and the geometric repetition of inputs across compartments leads the model to learn the average predictive structure across all compartments rather than the predictive structure particular to each compartment. In sum, among models tested, only a model with geometrically structured inputs and predictive encoding that is learned separately for each compartment can account for both individual differences in PV structure and the correlation between PV structure and successor structure that we observe in our actual data. For completeness, we provide additional visualizations of output from this model (Fig. 6d,e,f).

## Discussion

Here we leveraged miniscope imaging of mouse CA1 in a multicompartment environment to test whether patterns of remapping exhibited hallmarks of predictive structure in a complex environment. We showed that different mice instantiated different patterns of remapping across identically shaped compartments in this paradigm. These patterns coincided with the similarity of predictive navigational structure on a seconds-long timescale, modestly overweighting early experience. Limiting navigational options in this environment induced a new pattern of remapping which specifically matched the new predictive structure on this spatiotemporal scale. Finally, through computational modeling, we show that key results can be account for by a specific combination of predictive encoding over geometrically structured inputs learned separately within each compartment. Together, these results demonstrate that the structure of the hippocampal representation in a complex environment can be predicted from the behavioral history of the navigator, consistent with a predictive theory of cognitive mapping.

In this experiment, we compared patterns of remapping across identical unrewarded compartments from naiveté through extend experience to isolate a potential influence of predictive navigational structure on the hippocampal representation distinct from confounding differences in world features, reward structure, or previous experience. Given that we find evidence of such a determinant under these conditions, we hypothesize that remapping across environments intentionally disambiguated by external cues^36,41,43,44^ reflects both differences in input features as well as idiosyncratic differences in navigation. Likewise, these results provide evidence that predictive structure is an ongoing determinant of CA1 representational structure even in the absence of external rewards. Thus, we hypothesize that influences of reward structure on hippocampal coding^60–62^ may reflect both differences in rewarding input features as well as differences in goal-directed navigational trajectories. Our findings also suggest that idiosyncratic predictive structure likely contributed to the results of other multicompartment paradigms^39,47,52,53^ even when individual differences in representational structure may have been masked by averaging across animals.

We relied on the successor representation as an operationalization of a predictive theory of hippocampal cognitive mapping^19^. In this model, the hippocampus can be understood as mapping temporally discounted transitions between states of the world, which can be defined not only by location but also extra-spatial features. Indeed, one of the most compelling successes of this model is the ability to make quantitative predictions across scales (single cells, neural populations, behavior), species (rodents, humans), and domains (physical space, visual spaces, conceptual spaces)^19,63–65^. Although for simplicity we characterize transitions between locations as the representational determinant in this work, we imagine that in practice the CA1 representational structure may be better understood as reflecting transitions among its spatially varying inputs, including those from CA3 and entorhinal cortex^24^. Our modeling reflects this. In line with this reasoning, we would hypothesize that predictive structure determines CA1 structure when navigating extra-spatial domains, in both rodents and humans. Additionally, we note that the successor representation is not the only theory consistent with a predictive determinant of CA1 representational structure^20–22,66,67^. While our results are consistent with the predictions of this model, our experiment was not designed to adjudicate between competing theories of predictive cognitive mapping.

Our results demonstrate that CA1 representational structure most closely matches predictive structure when computed with a seconds-long temporal discount factor and overweighting early sessions. These characteristics are reminiscent of a unique form of hippocampal plasticity known as behavioral timescale synaptic plasticity (BTSP)^68,69^. BTSP is a mechanism by which new place fields can be endogenously or artificially induced in previously silent CA1 cells by evoking a calcium-mediated plateau potential. Properties of induced place fields indicate that inputs active within a few seconds of this event are potentiated, that these fields exhibit skew which depends on the navigational trajectory during the initial plateau potential, and that endogenous plateau potentials are most common during initial experience^68,70^. It is thus possible that the predictive determinants of CA1 representational structure we characterize here are a product of BTSP, and more generally that BTSP provides a biological basis for the normative claims of predictive mapping theories. If so, then we would expect to observe an increase in plateau potentials not only in novel environments but also during novel experiences which provoke representational change^71^. Our observations of changes in sparsity within and across experimental phases are roughly consistent with this. However, we interpret this finding with caution due to limitations inherent in our recording technique which do not allow us to distinguish plateau potentials from typical action potentials.

The critical spatiotemporal scale of predictive structure which we characterize here may also depend upon recording location within the hippocampus. It is known that the scale of spatial representations differs along the long dorsoventral axis of the hippocampus, progressing from finely tuned cells in the dorsal hippocampus to coarsely-tuned cells in the ventral^72^. Entorhinal inputs mirror this arrangement^73–75^. Even if the timescale of plasticity which encodes predictive structure were to be the same across this axis, differences in the scale of inputs might produce differences in the scale of encoded predictive structure^76^. In our work here, all populations were recorded from the dorsal hippocampus. Future work recording from the ventral hippocampus under similar conditions might address this possibility.

## Methods

### Subjects

Eleven naive mice (C57Bl/6, Charles River; 6 male, 5 female) were housed in pairs in 20 cm x 40 cm cages which included running wheels and additional enrichment. Mice were kept on a 14-hour light/10-hour dark cycle at 23°C and 30% humidity with food and water ad libitum. All experiments were carried out during the light portion of the light/dark cycle, and in accordance with University of Illinois Chicago Animal Use and Care Committee (protocol #23008) and with AAALAC guidelines.

### Surgeries

During all surgeries, mice were anesthetized via inhalation of a combination of oxygen and 5% Isoflurane before being transferred to the stereotaxic frame (David Kopf Instruments), where anesthesia was maintained via inhalation of oxygen and 0.5-2.5% Isoflurane for the duration of the surgery. Body temperature was maintained with a heating pad and eyes were hydrated with gel (Optixcare). Meloxicam (2 mg kg^-1^) and saline (0.5 ml) were administered subcutaneously at the beginning of each surgery. Preparation for recordings involved three surgeries per mouse.

First, at the age of six to ten weeks, each mouse was transfected with a 400 nl injection of the calcium reporter GCaMP8f via the viral construct AAV9.syn.GCaMP8f.WPRE.SV40 with an original titre of 2.3 x 10^13^ GC ml^-1^ (Addgene) diluted at a 1 part virus to 7 parts sterile artificial cerebrospinal fluid before surgical microinjection. While expression in both principal cells and inhibitory populations is possible under the syn promoter, it is likely that the large majority of cells we record are excitatory given their sparse transients.

Three weeks post-injection, a 1.8mm diameter gradient refractive index (GRIN) lens (Edmund Optics) was implanted above dorsal CA1 (Referenced to bregma: ML = 2.0 mm, AP = - 2.1 mm; Referenced to brain surface: DV = -1.35 mm). Implantation required aspiration of intervening cortical tissue. In addition to the GRIN lens, two stainless steel screws were threaded into the skull above the contralateral hippocampus and prefrontal cortex to stabilize the implant. Dental cement (C&B Metabond) was applied to secure the GRIN lens and anchor screws to the skull. A silicone adhesive (Kwik-Sil, World Precision Instruments) was applied to protect the top surface of the GRIN lens until the next surgery.

Three weeks after lens implantation, an aluminum baseplate was affixed via dental cement (C&B Metabond) to the skull of the mouse, which would later secure the miniaturized fluorescent endoscope (miniscope) in place during recording. The miniscope/baseplate was mounted to a stereotaxic arm for lowering above the implanted GRIN lens until the field of view contained visible cell segments and dental cement was applied to affix the baseplate to the skull. A polyoxymethylene cap was affixed to the baseplate when the mice were not being recorded to protect the baseplate and lens.

After surgery, animals were continuously monitored until they recovered. For the initial two days after surgery mice were provided with additional doses of meloxicam for pain management. One week following baseplating, to familiarize mice with the recording procedure and to monitor the quality of cell activity, we recorded mice daily in a 75 cm x 75 cm square open field for 10 min to 30 min per day. When recording quality was deemed of sufficiently high quality based on a stable number of cells across days, rapid transients, and a high proportion of cells with stable place fields within day (>50% of cells versus shuffled controls), mice began the multicompartment experiment. Mice typically met these criteria 2+ weeks following baseplating.

### Apparatus

The recording environment was constructed of white Lego base and white acrylic walls (Professional Plastics). All walls had a height of 20 cm. All compartments were 20 cm x 35 cm, with a 20cm central chamber. No other internal directional cues were provided, but the apparatus was closer to one external grey room wall which could serve as an external directional cue. During recording, the environment was dimly lit by an LED lamp positioned to reduce shadows, and a white-noise machine was used to mask any uncontrolled sounds. All sessions were 30 or 40 min (depending on signal strength to minimize photobleaching), and only one session was recorded per day. On rare occasions (∼2%) sessions were impacted by equipment failures, and sessions were terminated early; data from these sessions were not analyzed. During all free exploration sessions, mice were placed in the center of the environment at the start of the session facing the nearer external grey room wall. During limited navigation sessions, mice were placed in the first room at the start of the session also facing the nearer external grey wall. Mice were not intentionally disoriented prior to recording, and the stability of patterns of remapping across days suggests that they remained oriented throughout. The recording environment was cleaned between recordings with veterinarian-grade disinfectant.

### Data acquisition

In vivo calcium videos were recorded with a UCLA miniscope (v3; miniscope.org) containing a monochrome CMOS imaging sensor (MT9V032C12STM, ON Semiconductor) connected to a custom data acquisition (DAQ) box (miniscope.org) with a lightweight, flexible coaxial cable. The DAQ was connected to a PC with a USB 3.0 SuperSpeed cable and controlled with Miniscope custom acquisition software (miniscope.org; software version v4). The outgoing excitation LED was set to between 5-30%, depending on the mouse to maximize signal quality with the minimum possible excitation light to mitigate the risk of photobleaching. Gain was adjusted to match the dynamic range of the recorded video to the fluctuations of the calcium signal for each recording to avoid saturation. Behavioral video data were recorded by a webcam mounted above the environment. The DAQ simultaneously acquired behavioral and cellular imaging streams at 30 Hz as FFV1 losslessly-compressed AVI files and all recorded frames were timestamped for post-hoc alignment.

### Data preprocessing

Calcium imaging data were preprocessed prior to analyses via a pipeline of open source MATLAB (MathWorks; version R2024a) functions to correct for motion artifacts^77^, segment cells and extract transients^78^. The motion-corrected calcium imaging data were manually inspected to ensure that motion correction was effective and did not introduce additional artifacts. Following this preprocessing pipeline, the spatial footprints of all cells were manually verified to remove lens artifacts. Finally, to estimate the spike trains which gave rise to the transients we recorded, we implemented a second-order autoregressive deconvolution algorithm^51^. This nonnegative vector was then binarized such that any frame with a nonzero likelihood was treated as active (i.e. 1), otherwise it was treated as not active (i.e. 0). All further analysis was conducted on this binary activity vector.

Position data were inferred from behavioral videos via DeepLabCut^79^ tracking the nose, ears, and tail base of the mouse. The average between the two ears was taken as the position of the mouse in all analyses. Periods were estimates of either ear positions were low-confidence (<0.90) or the estimated distance between the ears was unrealistic were excluded and position was linearly interpolated across these timepoints (<0.5% of timepoints). Position data were then resampled via linear interpolation based on system clock timestamping to estimate the position at each time when imaging frames were collected.

### Data analysis

All analyses were conducted using the binarized nonnegative spike train estimates inferred via a second-order autoregressive deconvolution, henceforth the *activity* vector.

#### Rate maps

Rate maps for each cell were computed by dividing the environment (or each compartment) into 2.5 cm x 2.5 cm pixels, computing the mean activity rate at each pixel for that cell, and then smoothing the result map with an isotropic gaussian kernel with standard deviation of 2.5 cm. These maps were then corrected for edge artifacts and artifacts induced by smoothing over unsampled pixels as follows (MATLAB function *nanconv* available at https://www.mathworks.com/matlabcentral/fileexchange/41961-nanconv). First, a second map where sampled pixels are set to one and all other pixels are set to zero was computed. This maps was then smoothed with the same kernel, and each rate map was divided by this smoothed map. Unsampled pixels were excluded from further comparisons.

#### Split-half stability

To characterize the stability of each cell’s spatial tuning during a session, we computed its split-half stability by computing whole environment rate maps using only the first half or second half of the data from that session and correlating the two rate maps. To determine the statistical significance of these correlations, we compared this true correlation to a surrogate distribution computed by randomly circularly shifting the position vector relative to the activity vector by at least 30 seconds and recomputing the resulting correlation 1000 times. Cells with true correlations exceeding the 95% of this surrogate distribution were considered spatially-stable and included in further analysis.

#### Spatial information content

To characterize the specificity of each cell’s spatial tuning during a session, we computed its spatial information content from its whole environment rate map as described previously^80^ via the equation:

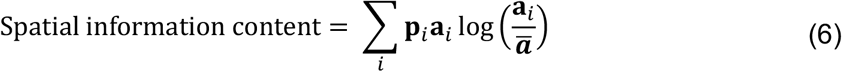

where *i* is the rate map pixel index, **p**_*i*_ is the probability of sampling pixel *i*, **a**_*i*_ is the mean firing rate at pixel *i*, and **ā** is the mean firing rate across all pixels. To determine the statistical significance of these values, we compared the true spatial information content to a surrogate distribution computed by randomly circularly shifting the position vector relative to the activity vector by at least 30 seconds and recomputing the resulting correlation 1000 times. Cells with true correlations exceeding the 95% of this surrogate distribution were considered spatially tuned and included in further analysis.

#### Similarity structures

Population vector correlations between compartments were computed by aligning these rate maps by the entryway, concatenating the linearized compartment rate maps of all cells from each compartment, and computing the correlation between these vectors. This measure of takes into account both changes in the spatial distributions of firing and relative changes in firing rate. Mean activity rate correlations between compartments were computed by correlating the vector of mean activity rates for all included cells between compartments. Successor structure and cumulative successor structure were computed as described in the main text. To compute successor matrices, position data were binned to 5 cm x 5 cm pixels, and the timestep was set to 1/30^th^ of a second to match the frame rate of data acquisition.

### Computational modeling

To compute activity vectors of model inputs, first each compartment was divided into a grid of possible preferred locations at 1.25 cm x 1.25 cm increments (448 possible preferred locations). Each cell was assigned one of these preferred locations (448 input cells total). For geometrically structured inputs, each cell had the same preferred location across all compartments, aligned by their entryways. For randomly structured inputs, the assignment of preferred locations was randomly shuffled across cells for each compartment. Once assigned, each cell kept its preferred locations for all sessions. At each timestep, the activity of each cell was determined from its distance to its nearest preferred location according to a gaussian distribution with standard deviation 5 cm. Finally, input activity vectors were normalized to scale from [0, 1] inclusive. Each transformation received identical input traces for the geometrically structured inputs and randomly structured inputs.

For the linear transformation, each CA1 cell received a random number of inputs on the range [2, 16] drawn from a Poisson distribution with *λ* = 4. These inputs were chosen from the input population at random and given equal weight (i.e. 1). All other inputs were assigned a weight of 0 for that CA1 cell. Once assigned, these weights were constant across all sessions. SR and multi-SR transformations are described in the main text. In all cases, the successor matrices were initialized as the identity matrix.

Finally in all cases the output *O*_*i*_ from CA1 cell *i* was determined at each timestep by

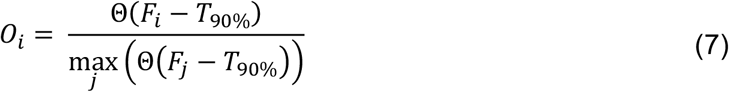

where *F*_*i*_ is the total input to CA1 cell *i, T*_90%_ is a dynamic threshold corresponding to the *F* of the cell with the 90^th^ percentile *F* across the population at that timestep, and

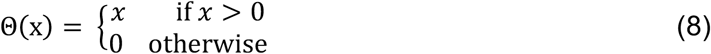

The result of this thresholding and normalization is that only 10% of CA1 cells are active during each timestep with output rates ranging from [0, 1]. This dynamic normalization is intended to mimic competition among principal cells in CA1^81^, and is especially useful when changes in connectivity during learning lead to large changes in excitatory drive as is often the case in our SR and multi-SR simulations. For each model, we simulated 448 CA1 cells, matching the number of input cells. Output vectors were treated like the activity vectors of our actual data in all further analyses. SR and multi-SR transformations were deterministic and therefore a single iteration is presented in the main text. A single iteration of the linear transformation model was also included to match the presentation of SR and multi-SR simulations, despite some additional randomness in its implementation. However, we note that we have run approximately 10 iterations at the identical parameterization and in all cases the results closely resemble those presented in the main text.

### Histological validation of expression and recording targets

After experiments, animals were perfused to verify GRIN lens placement. Mice were deeply anesthetized and intracardially perfused with 4% paraformaldehyde in PBS. Brains were dissected and post-fixed with the same fixative. Coronal sections (50 μm) of the entire hippocampus were cut using a vibratome and sections were mounted directly on glass slides. Sections were split and half of all sections were stained for DAPI and mounted with Fluoromount-G (Southern Biotechnology) to localize GRIN lens placement and to evaluate viral expression. Due to the large imageable surface but restricted miniscope field of view (∼0.5 mm x ∼0.8 mm), we were unable to determine more specific localization of populations within the hippocampus for mice recorded with 1.8 mm lenses.

### Statistics and reproducibility

All statistical tests are noted where the corresponding results are reported throughout the main text and supplement. All tests were uncorrected 2-tailed tests unless otherwise noted. Z-values for nonparametric rank-sum and signed-rank tests were not estimated or reported for comparisons with fewer than 15 datapoints. Box plots portray the minimum and maximum (whiskers), upper and lower quartiles (boxes), and median (cinch/bolded line). All correlations are Pearson’s correlations unless otherwise noted.

### Code availability

All custom code written for reported analyses are publicly available at [insert Github link] or via request to the corresponding authors.

### Data availability

The complete dataset for all experiments are publicly available at [insert Dryad link] or via request to the corresponding authors.

## Acknowledgements

This work was supported by startup funding provided by the University of Illinois Chicago, as well as a UIC LAS CSSR seed grant. We would also like to thank J. Quinn Lee, Rachel Donka, and Rachel Barrett for feedback when drafting this manuscript.

## Author Contributions

SAP and ATK contributed to experimental design, surgeries, recordings, analysis of data, as well as revising the manuscript. ATK drafted the manuscript.

## Supplementary information for

**Supplementary Figure 1.**
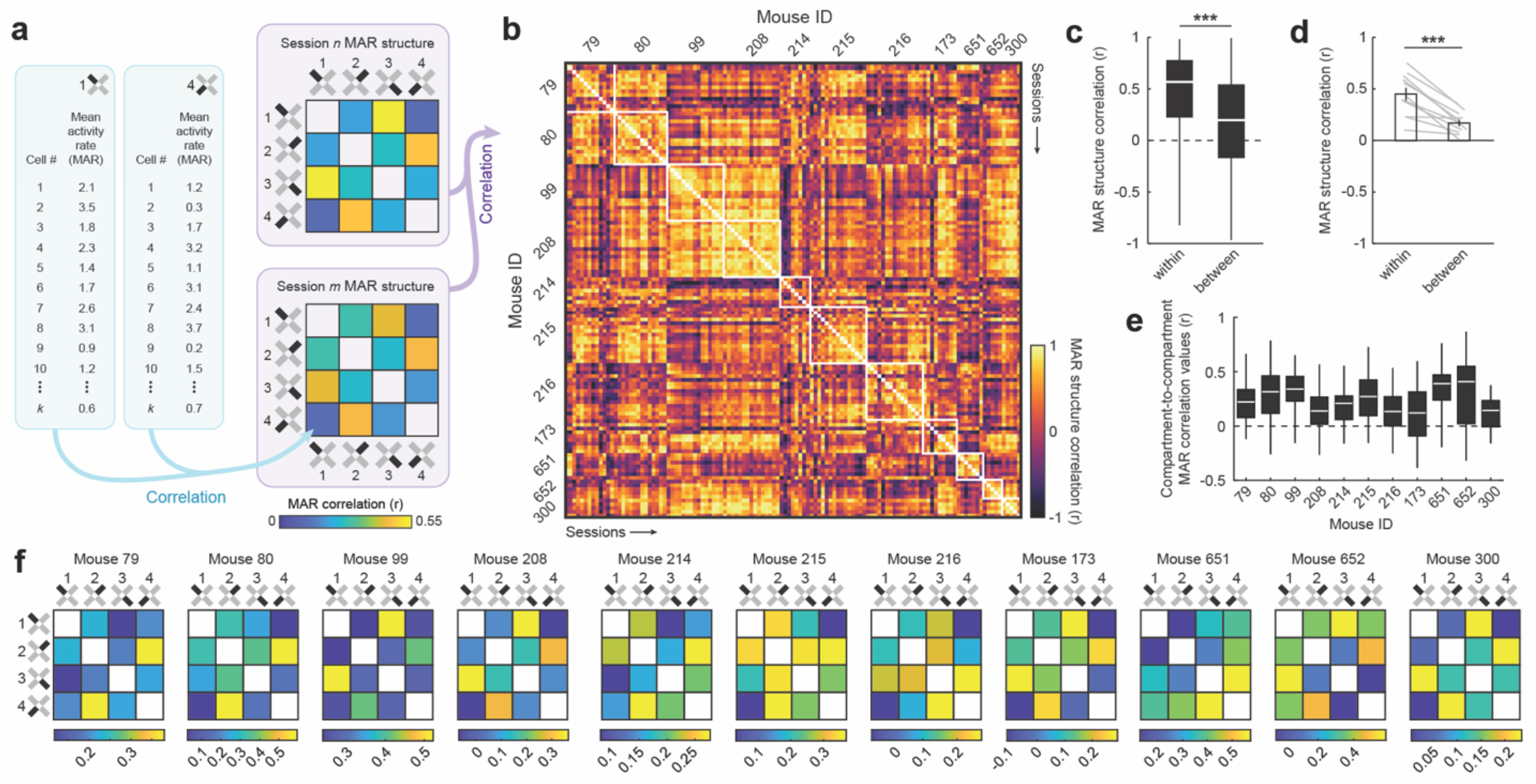
Individual differences in patterns of remapping across compartments in a multicompartment environment as assessed by comparing mean activity rate. (a) Schematic of mean activity rate (MAR) structure analysis pipeline. (b) MAR structure correlations across all pairwise comparisons of sessions. White boxes bound session comparisons from the same mice. (c) MAR structure correlations for comparisons within mouse versus between different mice, treating each pairwise session comparison independently (rank-sum test: z(3395800) = 16.687, p = 16.687). (d) MAR structure correlations for comparisons within mouse versus between mice, treating each mouse as the unit of analysis (signed-rank test: p = 9.766e-04). Soft lines denote individual mice. (e) Distribution of compartment-to-compartment MAR correlation values for each mouse. (f) MAR structures averaged across sessions for each mouse. ***p<0.001

**Supplementary Figure 2.**
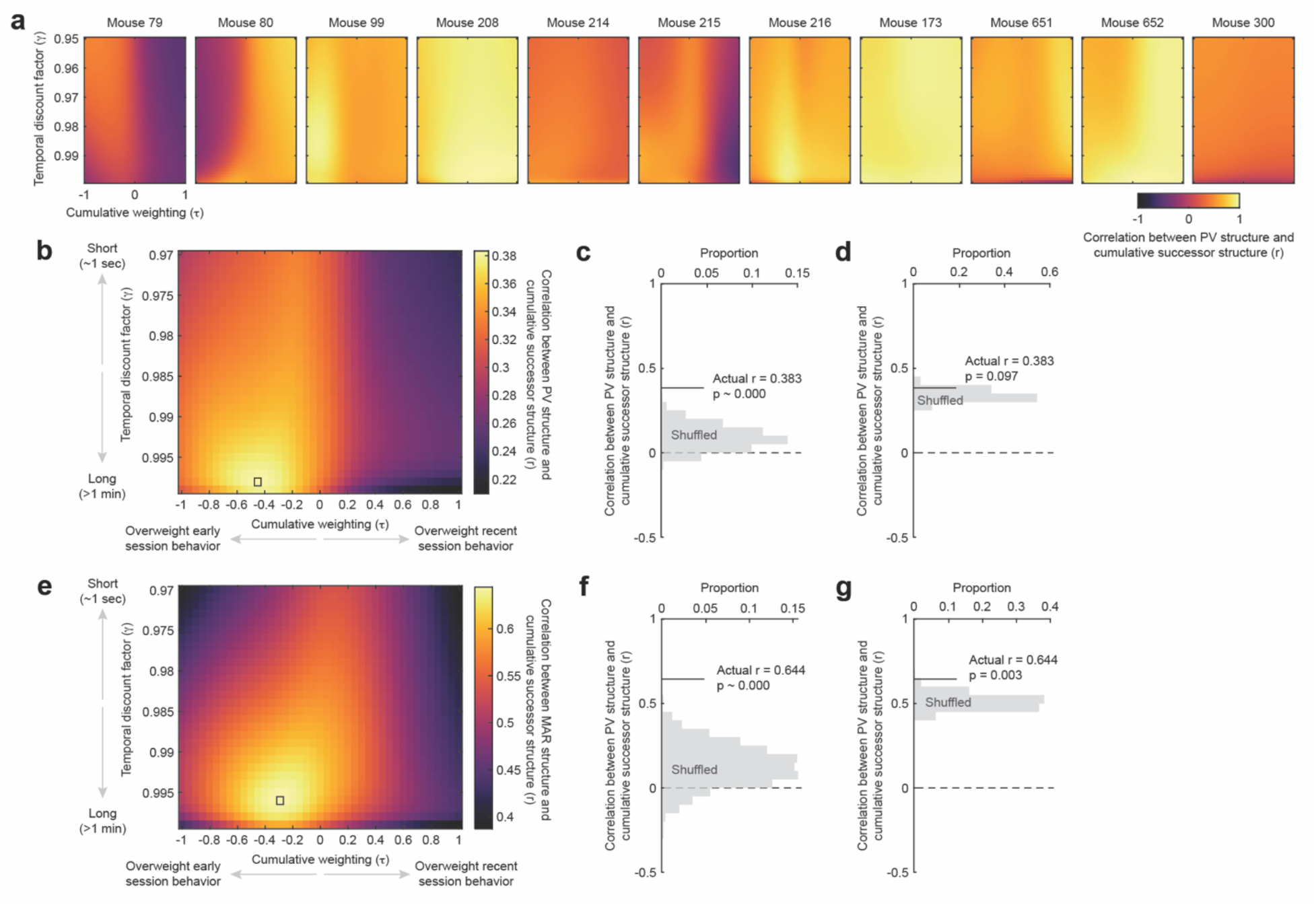
Patterns of remapping in CA1 resemble the similarity of predictive navigational structure on a particular spatiotemporal scale. (a) Grid search where γ and τ are varied and the mean cumulative successor structure is correlated with the mean PV structure shown separately for each mouse. (b) Grid search where γ and τ are varied and the cumulative successor structure is correlated with the PV structure, treating each session as an independent statistical unit. (c) Correlation between PV structure and cumulative successor structure against a shuffled control where transition probabilities are shuffled across compartments, mice, and sessions and controlling for the grid search, treating each session as an independent statistical unit. (d) Correlation between PV structure and cumulative successor structure against a shuffled control where transition probabilities are shuffled across sessions within mouse and controlling for the grid search, treating each session as an independent statistical unit. (e) Grid search where γ and τ are varied and the mean cumulative successor structure is correlated with the mean activity rate (MAR) structure for each mouse. (f) Correlation between MAR structure and cumulative successor structure against a shuffled control where transition probabilities are shuffled across compartments, mice, and sessions and controlling for the grid search, averaging within mouse prior to correlation. (g) Correlation between PV structure and cumulative successor structure against a shuffled control where transition probabilities are shuffled across sessions within mouse and controlling for the grid search, averaging within mouse prior to correlation.

**Supplementary Figure 3.**
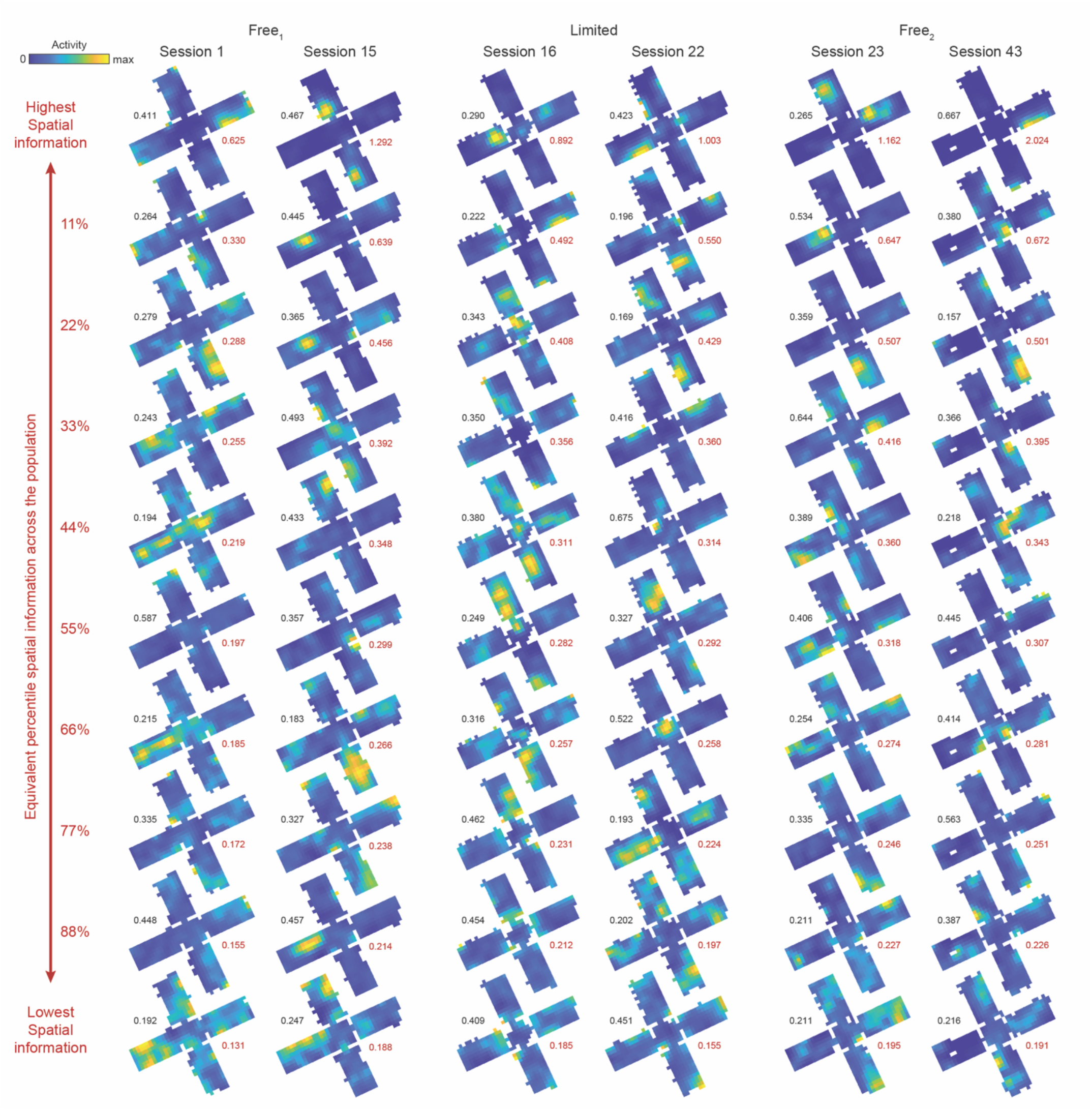
Example rate maps for all experimental phases. Example rate maps for the first and last session of each phase for Mouse 208. Examples chosen from the equivalent percentile of the spatial information distribution for each session, ordered from highest to lowest. Peak activity rate noted for each rate map in black. Spatial information noted for each rate map in red.

**Supplementary Figure 4.**
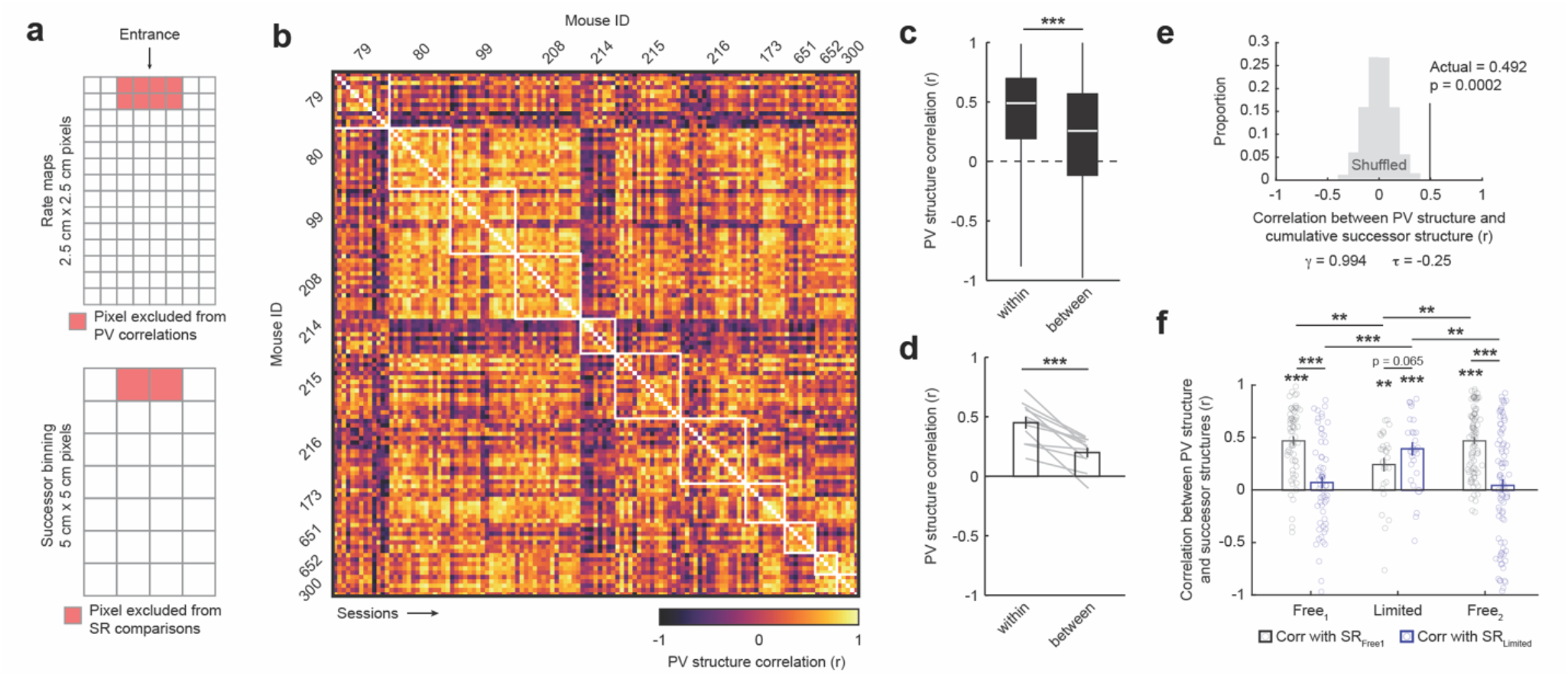
Excluding data recorded near the entryway did not substantially change the results. (a) For all analyses in this figure, data from the pixels near the entryway were excluded from both the PV structure and successor structure computations. (b) PV structure correlations across all pairwise comparisons of sessions when excluding entry pixels. White boxes bound session comparisons from the same mice. (c) PV structure correlations for comparisons within mouse versus between different mice when excluding entry pixels, treating each pairwise session comparison independently (rank-sum test: z(3148018) = 11.968, p = 5.207e-33). (d) PV structure correlations for comparisons within mouse versus between mice when excluding entry pixels, treating each mouse as the unit of analysis (signed-rank test: p = 9.766e-04). Soft lines denote individual mice. (e) Mean correlation between PV structure and cumulative successor structure at the parameterization γ = 0.994, τ = -0.25 when excluding entry pixels. Shuffled distribution is the mean correlation across 10,000 random shuffles of transition matrices across compartments, sessions, and mice prior to computing cumulative successor structures. P-value estimated nonparametrically from this distribution. (f) Correlation between PV structure and both SR_Free1_ and SR_Limited_ for all sessions, aggregating across mice. For complete statistical information, see Supplementary Table 1. **p<0.01, ***p<0.001

**Supplementary Table 1.** Statistical outcomes for comparisons of correlation between PV structures and both SR_Free1_ and SR_Limited_ as in Figure 4c and Supplementary Figure 3.

| Experimental Phase | Comparisons | Statistical test | Statistic | Outcome |
| --- | --- | --- | --- | --- |
| Main text results (Figure 4c) |  |  |  |  |
| Free<sub>1</sub> | SR<sub>Free1</sub> vs. 0 | Signed rank | 1762 | 4.510e-10 |
| Free<sub>1</sub> | SR<sub>Limited</sub> vs. 0 | Signed rank | 1151 | 8.233e-02 |
| Free<sub>1</sub> | SR<sub>Free1</sub> vs. SR<sub>Limited</sub> | Signed rank | 1528 | 6.402e-06 |
| Limited | SR<sub>Free1</sub> vs. 0 | Signed rank | 346 | 1.129e-03 |
| Limited | SR<sub>Limited</sub> vs. 0 | Signed rank | 363 | 2.690e-04 |
| Limited | SR<sub>Free1</sub> vs. SR<sub>Limited</sub> | Signed rank | 124 | 7.203e-02 |
| Free<sub>2</sub> | SR<sub>Free1</sub> vs. 0 | Signed rank | 3484 | 3.532e-14 |
| Free<sub>2</sub> | SR<sub>Limited</sub> vs. 0 | Signed rank | 1942 | 4.838e-01 |
| Free<sub>2</sub> | SR<sub>Free1</sub> vs. SR<sub>Limited</sub> | Signed rank | 3272 | 3.319e-11 |
| Free<sub>1</sub> vs. Limited | SR<sub>Free1</sub> | Rank sum | 3064 | 4.231e-04 |
| Free<sub>1</sub> vs. Limited | SR<sub>Limited</sub> | Rank sum | 2385 | 1.081e-02 |
| Free<sub>1</sub> vs. Free<sub>2</sub> | SR<sub>Free1</sub> | Rank sum | 4263 | 7.260e-01 |
| Free<sub>1</sub> vs. Free<sub>2</sub> | SR<sub>Limited</sub> | Rank sum | 4431 | 7.443e-01 |
| Limited vs. Free<sub>2</sub> | SR<sub>Free1</sub> | Rank sum | 1011 | 1.264e-04 |
| Limited vs. Free<sub>2</sub> | SR<sub>Limited</sub> | Rank sum | 1987 | 6.568e-03 |
| With pixels near the entrance removed (Supplementary Figure 3) |  |  |  |  |
| Free<sub>1</sub> | SR<sub>Free1</sub> vs. 0 | Signed rank | 1771 | 2.947e-10 |
| Free<sub>1</sub> | SR<sub>Limited</sub> vs. 0 | Signed rank | 1088 | 2.028e-01 |
| Free<sub>1</sub> | SR<sub>Free1</sub> vs. SR<sub>Limited</sub> | Signed rank | 1553 | p = 2.644e-06 |
| Limited | SR<sub>Free1</sub> vs. 0 | Signed rank | 344 | 1.324e-03 |
| Limited | SR<sub>Limited</sub> vs. 0 | Signed rank | 379 | 6.129e-05 |
| Limited | SR<sub>Free1</sub> vs. SR<sub>Limited</sub> | Signed rank | 122 | 6.511e-02 |
| Free<sub>2</sub> | SR<sub>Free1</sub> vs. 0 | Signed rank | 3504 | 1.770e-14 |
| Free<sub>2</sub> | SR<sub>Limited</sub> vs. 0 | Signed rank | 1967 | 4.170e-01 |
| Free<sub>2</sub> | SR<sub>Free1</sub> vs. SR<sub>Limited</sub> | Signed rank | 3020 | 3.633e-08 |
| Free<sub>1</sub> vs. Limited | SR<sub>Free1</sub> | Rank sum | 3007 | 2.573e-03 |
| Free<sub>1</sub> vs. Limited | SR<sub>Limited</sub> | Rank sum | 2292 | 7.199e-04 |
| Free<sub>1</sub> vs. Free<sub>2</sub> | SR<sub>Free1</sub> | Rank sum | 4390 | 8.728e-01 |
| Free<sub>1</sub> vs. Free<sub>2</sub> | SR<sub>Limited</sub> | Rank sum | 4374 | 9.241e-01 |
| Limited vs. Free<sub>2</sub> | SR<sub>Free1</sub> | Rank sum | 1172 | 5.930e-03 |
| Limited vs. Free<sub>2</sub> | SR<sub>Limited</sub> | Rank sum | 2015 | 3.659e-03 |

